# Miniaturization and optimization of 384-well compatible metagenomic sequencing library preparation

**DOI:** 10.1101/440156

**Authors:** Madeline Y Mayday, Lillian M Khan, Eric D Chow, Matt S Zinter, Joseph L DeRisi

## Abstract

Preparation of high-quality sequencing libraries is a costly and time-consuming component of metagenomic next generation sequencing (mNGS). While the overall cost of sequencing has dropped significantly over recent years, the reagents needed to prepare sequencing samples are likely to become the dominant expense in the process. Furthermore, libraries prepared by hand are subject to human variability and needless waste due to limitations of manual pipetting volumes. Reduction of reaction volumes, combined with sub-microliter automated dispensing of reagents without consumable pipette tips, has the potential to provide significant advantages. Here, we describe the integration of several instruments, including the Labcyte Echo 525 acoustic liquid handler and the iSeq and NovaSeq Illumina sequencing platforms, to miniaturize and automate mNGS library preparation, significantly reducing the cost and the time required to prepare samples. Through the use of External RNA Controls Consortium (ERCC) spike-in RNAs, we demonstrated the fidelity of the miniaturized preparation to be equivalent to full volume reactions. Furthermore, detection of viral and microbial species from cell culture and patient samples was also maintained in the miniaturized libraries. For 384-well mNGS library preparations, we achieved a savings of over 80% in materials and reagents alone, and reduced preparation time by 90% compared to manual approaches, without compromising quality or representation within the library.

## Introduction

Metagenomic next-generation sequencing (mNGS) is becoming an increasingly useful tool in the field of biology and clinical medicine. Its applications are almost limitless – any nucleic acid can be turned into a library, amplified, and sequenced, making mNGS an appealing technology for labs and hospitals alike. As sequencers such as the Illumina NovaSeq increase throughput, hundreds to thousands of libraries can be sequenced in a single run. Although the per-base cost of sequencing has become less expensive over the last several decades, the cost and time associated with sample preparation remain disproportionately high [1,2].

Manual library preparation is tedious and is often the bottleneck for many sequencing projects. Numerous library preparation protocols have been adapted for automation through the use of various positive displacement tip-based liquid handler instruments, including the Beckman Coulter Biomek, Hamilton Star, Agilent Technologies Bravo, TTP LabTech Mosquito, and others [3–5]. Though these provide more hands-off time during the library preparation process, the overall cost can often exceed that of hand-prepared libraries due to the increased dead volume of reagents and the large number of expensive, sometimes proprietary tips required for liquid handlers. Furthermore, sub-microliter miniaturization is a challenge for the majority of positive displacement based liquid handlers.

Recently, acoustic liquid handlers with sub-microliter precision have become commercially available and have been used for a large range of applications, including RT-qPCR, mass spectrometry, drug discovery, and compound dosing assays [6–9]. To date, few end-to-end protocols for miniaturized mNGS preparation have been available. Given the cost and time limitations of current library preparation techniques, we sought to adapt our mNGS library preparation protocol into a high-throughput protocol by leveraging the small dispensing volumes of the Echo 525. Here, we describe a detailed protocol that provides high-fidelity, miniaturized, automated, cost- and time-efficient 384-well library preparation together with its quality and performance metrics.

## Methods and results

All next generation sequencing workflows begin with isolation of nucleic acid; however, such protocols are highly dependent on the source of the sample and desired product, such as DNA, RNA, mRNA, cell-free RNA, and so on. The protocol described herein is independent of nucleic acid isolation methods, and for the purpose of this work, we assume an input of isolated total RNA. To optimize miniaturization of our laboratory’s current library preparation protocol, we prepared libraries from varying concentrations of HeLa RNA using the New England Biolabs Ultra II Library Prep Kit (E7770S/L). The miniaturized and automated workflow is presented in **Fig 1** and described in detail below and at http://doi.org/10.17504/protocols.io.tcaeise.

**Fig 1.**
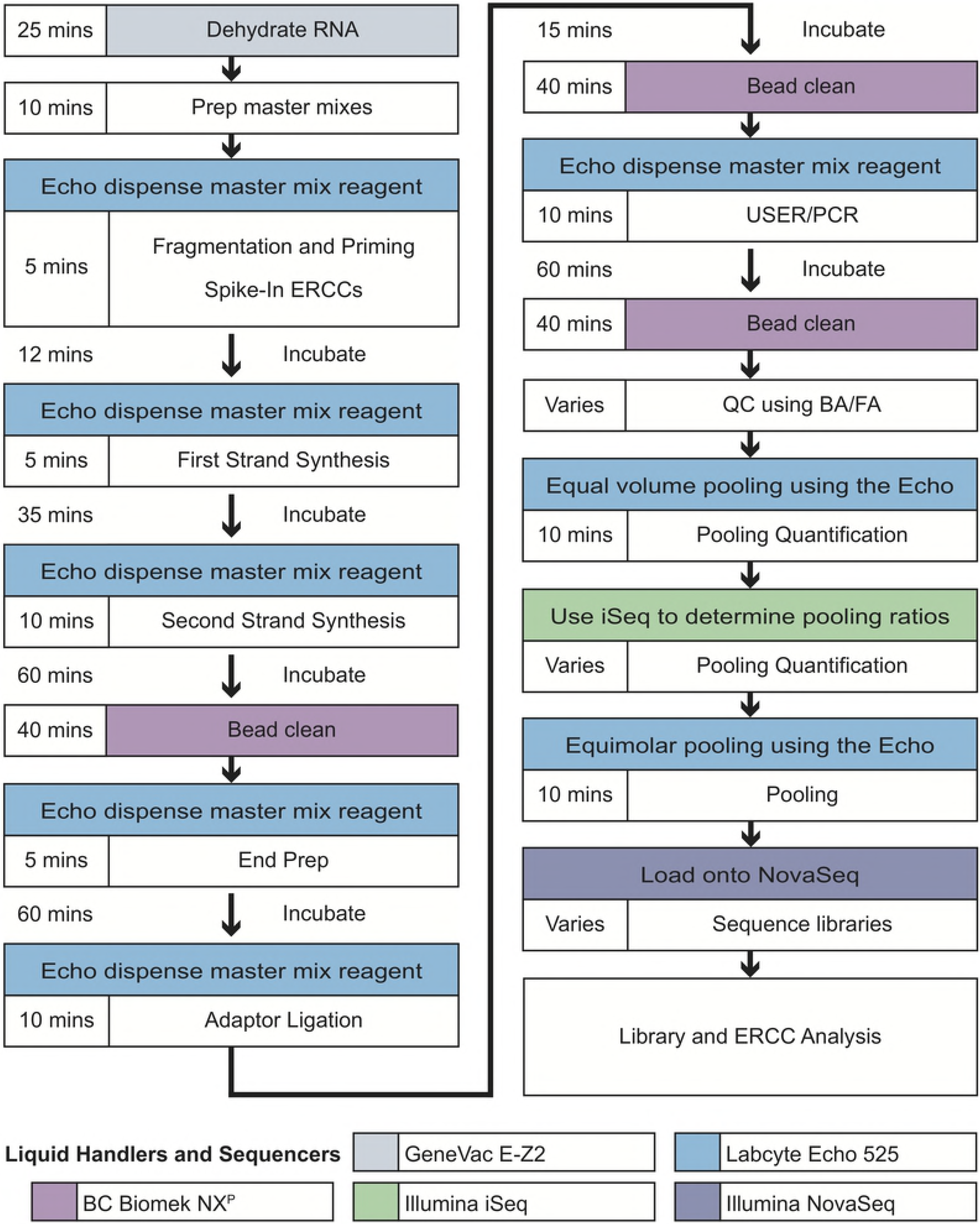
Overall workflow. **Legend:** Workflow of the miniaturized, automated library preparation protocol. The complete protocol is available at: http://doi.org/10.17504/protocols.io.tcaeise.

### Sample dehydration

To achieve miniaturized volumes, nucleic acid samples must first be dehydrated. Several vacuum evaporators were tested to dehydrate input RNA: Thermo Savant AES2010, Thermo Savant SC110A, Thermo Savant DNA110, and GeneVac EZ-2. 5uL of extracted RNA were loaded into a 384-well PCR plate (E&K Cat No EK-75009) and were spun at low (no heat/room temperature), medium (40°C), or high (60°C) temperatures until all wells were completely dry. Room temperature drying was prohibitively slow; at medium and high temperatures, drying times were comparable. At 40°C, sample plates dried fastest with the GeneVac E-Z2 (25 minutes) and ranged from 35-50 minutes using other machines. Of note, variable drying times were observed between brands of 384-well PCR plates. After drying RNA at 40°C for 25 minutes, samples were rehydrated in 5uL water. Parallel capillary electrophoresis (Agilent Bioanalyzer) RNA Integrity Number showed no significant difference between the resuspended sample and the original sample (T-test p=0.306, **Fig 2**), demonstrating that dehydration does not compromise RNA quality within these parameters.

**Fig 2.**
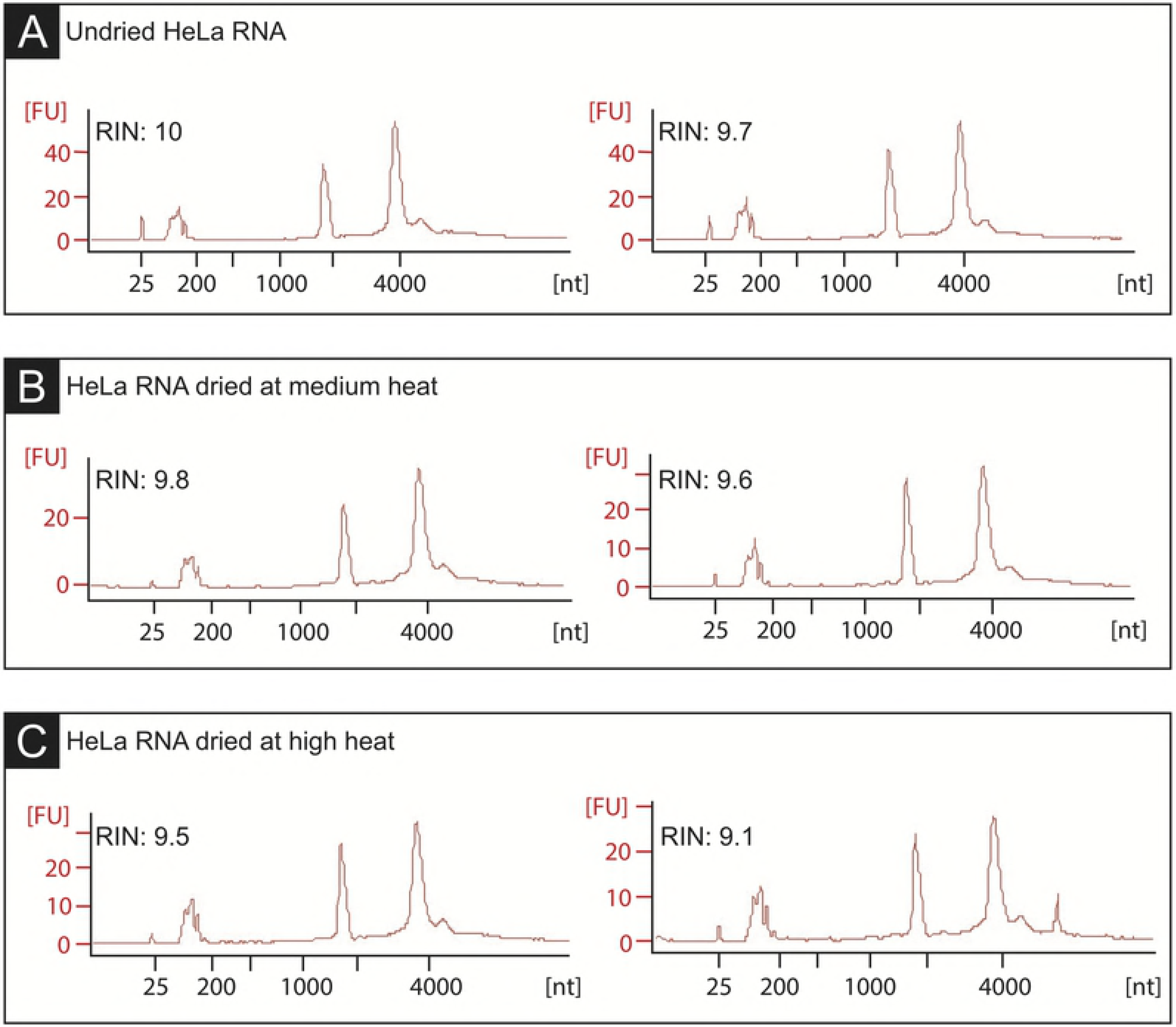
Dehydrated RNA demonstrates preserved integrity. **Legend:** Bioanalyzer traces and RNA Integrity Numbers (RINs) of biological replicates of HeLa RNA. (A) Before being dried in a vacuum evaporator. (B) After being dried for 30 minutes at 40°C. (C) After being dried for 25 minutes at 65°C. RINs indicate that RNA quality is not compromised during the dehydration process.

### Intra-sample positive control ERCC spike-ins

Internal spike-in control nucleic acids are useful indicators of potential library preparation errors. Furthermore, carefully designed spike-in controls, such as the External RNA Controls Consortium (ERCC) collection [10–12], which consists of 92 variable-length archaeal templates present at a pre-defined range of concentrations, may be used to establish the relationship between read count and input RNA concentration. For this mNGS protocol, 25 picograms of ERCC RNA (Thermo Fisher Scientific Cat. No 4456740) were added to each sample prior to library preparation.

### Library preparation

Our laboratory has previously described a mNGS library prep protocol for RNA using the New England Biolabs Ultra II Library Prep Kit [13–15]. In this protocol, RNA was quantified using QuBit, fragmented in a magnesium-based buffer at 94°C, primed with random hexamers, and reverse transcribed to form cDNA. Libraries were made Illumina-compatible by blunting ends of cDNA and adding non-templated d-A tails. Loop adaptors were ligated and cleaved with uracil-specific excision reagent (USER) enzyme before PCR enrichment was performed with sample-specific octamer primers.

To miniaturize and adapt this procedure to the Echo 525, we tested several protocol modifications involving reduced volume of reagents (**Table 1**) using variable HeLa RNA input (0, 0.1, 0.5, 1, and 5 ng in 5uL of water). Reagents for each step, including ERCC spike-ins, were prepared as a master mix and miniaturized volumes were dispensed using the Echo 525. The ideal number of PCR cycles is input-dependent and should be optimized depending on sample characteristics; for this miniaturized protocol, 19 PCR cycles were used to achieve adequate amplification of very low input samples.

**Table 1.** Library synthesis reactions at varying miniaturizations.

| Miniaturization | Sample input (uL) | RNA Fragmentation (uL) | First Strand Synthesis (uL) | Second Strand Synthesis (uL) | End Prep (uL) | Adaptor Ligation (uL) | USER/ PCR (uL) |
| --- | --- | --- | --- | --- | --- | --- | --- |
| **100% (full volume)** | 5 | 10 | 10 | 60 | 60 | 33.5 | 53 |
| **20%** | 1 | 2 | 2 | 12 | 12 | 6.7 | 15.9* |
| **12.5%** | 0.625 | 1.25 | 1.25 | 7.5 | 7.5 | 4.2 | 15.9* |
| **10%** | 0.5 | 1 | 1 | 6 | 6 | 3.3 | 15.9* |

**Legend:** Total volumes of master mixes for each step of the NEB Ultra II RNA Library Prep for each experimental miniaturization: 100%, 20%, 12.5% and 10% of the regular hand-prepped volume. *Maximum miniaturization was 30% volume in order to retain reaction efficacy.

Although all dispensing steps utilized acoustic liquid handling in this protocol, magnetic bead-cleaning steps still required pipetting automation. To facilitate high-throughput bead-cleaning, we programmed the Beckman Coulter Biomek NX^P^ with a 384 multichannel head to perform a simultaneous 384-well bead clean and size selection using AMPure XP beads (Beckman Coulter Cat. No A63881). Bead volumes were chosen to select for a final library size of≥200 base pairs and to remove adaptor- and primer-dimers. An Alpaqua 384 Post Magnet Plate (SKU A001222), adjusted with a 3D-printed plastic adaptor (available for download at http://derisilab.ucsf.edu/index.php?3D=225.), was used to minimize library elution volume. The adaptor raises the PCR plate when placed on the magnet plate, thereby lowering the height of the bead-DNA pellet on the side of each well and allowing for complete resuspension of beads in as low as 6uL of eluent.

To check the quality of libraries prepared using the miniaturized protocol, we used a parallel capillary electrophoresis assay to process up to 95 samples simultaneously (Advanced Analytical Fragment Analyzer). Libraries were assessed for distribution of cDNA fragment size, estimated molarity and concentration, and for the presence of primer- and adaptor-dimers. Library size distributions were consistent between libraries prepared with both full and miniaturized volumes (average fragment size=471±37bp, 86±7% between 200-1000bp).

### Sequencing results

Final HeLa RNA libraries were sequenced on the Illumina MiSeq to an average depth of 2.5 million paired-end reads. Resultant data were processed through a pipeline for pathogen detection developed in our laboratory involving subsequent removal of duplicate reads, reads with low quality, and reads aligning to phage [13,16,17]. Original FASTQ files are available at BioProject Accession #PRJNA493096.

Analysis of reads aligning to ERCC transcripts showed strong correlation to their original molar spike-in concentrations, which spanned six orders of magnitude, indicative of a successful library preparation and reflected uniform PCR amplification across fragment size (R^2^=0.93, **Fig 3A**). The linear association between input RNA concentration and sequencing output results was used to calculate the approximate amount of HeLa RNA present in the original sample by solving the following equation: 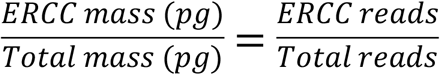. The mass of input RNA measured by fluorometric quantification (Qubit HS RNA) correlated strongly with estimations using the ERCC back-calculation method (R^2^=0.995, **Fig 3B**) indicating that ERCC controls are an effective way to assess the mass of input nucleic acid, even when present at sub-nanogram levels.

**Fig. 3.**
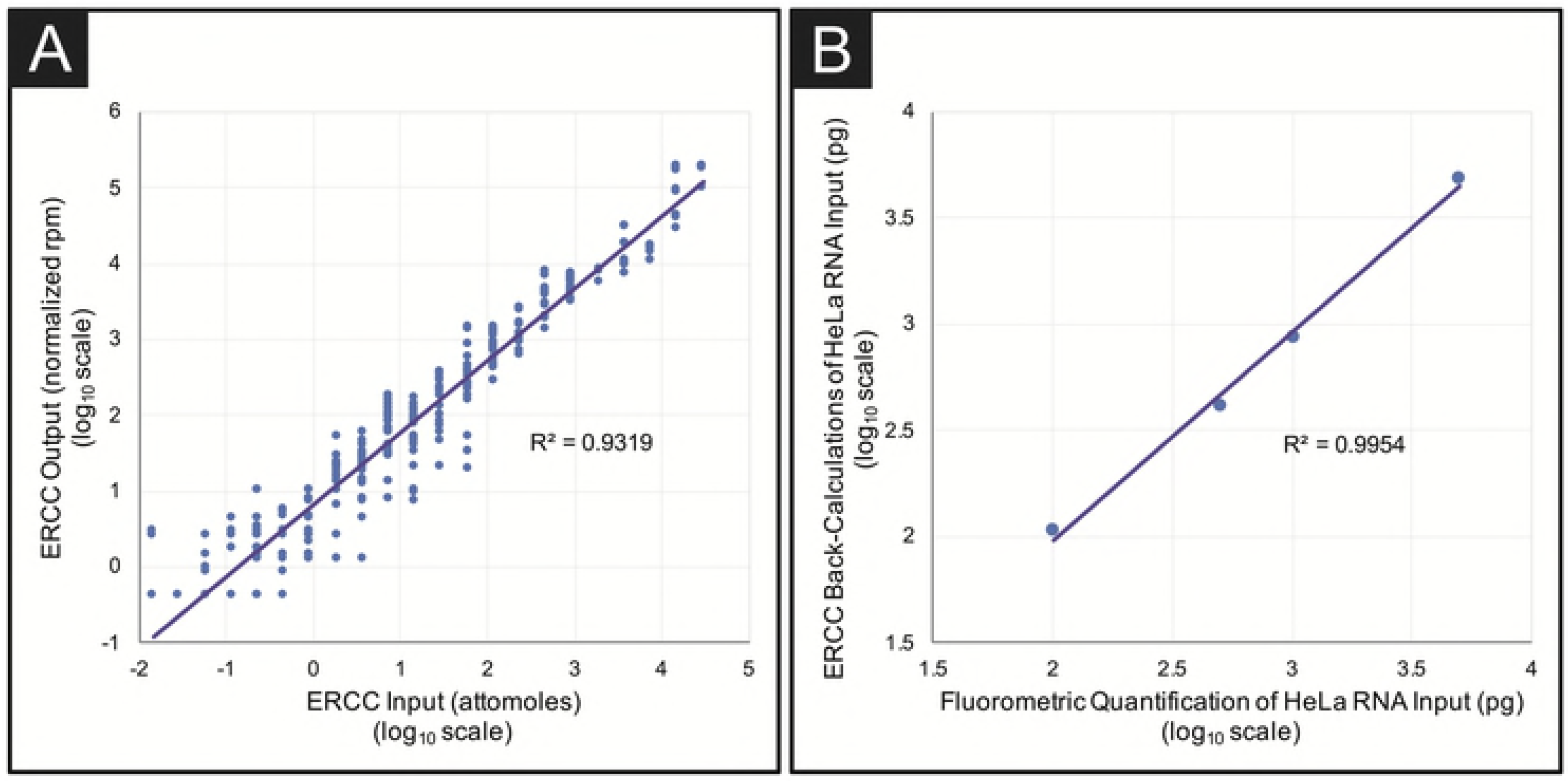
ERCC reads to determine library preparation quality and back-calculate RNA input mass. **Legend:** (A) After normalizing to RNA input mass, reads aligning to the 92 ERCC spike-in transcripts correlate linearly with ERCC spike-in concentration across six orders of magnitude in all libraries prepared with the miniaturized protocol (R^2^=0.932). (B) The initial sample input mass can be calculated using the ratio of ERCC reads to total sequencing reads in each sample using the equation: 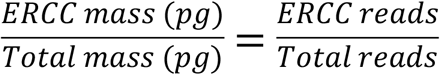 Back-calculated masses of HeLa libraries correlated strongly with QuBit quantification (R^2^=0.995).

The integrated portion of the HPV18 genome that is inherent to HeLa cells was used as a proxy for a human infection as would be expected in metagenomic sequencing of patient samples [18, 19]. Libraries were analyzed for percent and depth of coverage of the reference sequence of integrated HPV18, as well as the human transcriptome. At 1ng and 5ng of input, HPV18 coverage was essentially indistinguishable for reactions performed with full or miniaturized volumes of reagents, indicating that using as low as 10% of the standard reagent volume provided equivalent detection of the viral genome (**Table 2**). Assessment of human transcriptome coverage in the HeLa libraries demonstrated high correlation between full volume and miniaturized reactions (for 5ng RNA input: Spearman’s ρ = 0.79, p<0.001; for 1ng RNA input: Spearman’s ρ = 0.69, p<0.001, **Fig 4**). Additionally, the miniaturized protocol demonstrated strong correlation between replicate 1ng HeLa RNA samples (Spearman’s ρ = 0.84, p<0.001). Together, these metrics show that libraries prepared using the miniaturized protocol are equally sensitive for detection of both pathogen and human reads.

**Table 2).** HPV coverage in HeLa samples.

| HeLa RNA | Full (F)<br>Miniaturized (M) | Unique<br>paired-end<br>reads | Average depth | Integrated<br>HPV Coverage |
| --- | --- | --- | --- | --- |
| 5ng | F | 1,609,470 | 23.6±10.9 | 99.6% |
| 1ng | F | 930,857 | 8.0±5.0 | 91.5% |
| 5ng | M | 2,770,339 | 23.0±15.9 | 98.9% |
| 1ng (rep) | M | 2,597,928 | 17.5±9.7 | 100% |
| 1ng (rep) | M | 31,114,923 | 59.4±34.8 | 100% |
| 0.5ng | M | 2,427,233 | 14.3±10.0 | 99.5% |
| 0.1ng | M | 1,642,888 | 7.6±5.0 | 96.9% |

**Legend:** Coverage of the HPV genome was similar for libraries prepped with both the regular and miniaturized protocols with various RNA input concentrations. This indicates that pathogen detection is maintained despite the reduction in reagent volume. Miniaturized libraries were prepared using 10% of the original volumes, with the exception of PCR, as described above.

**Fig 4.**
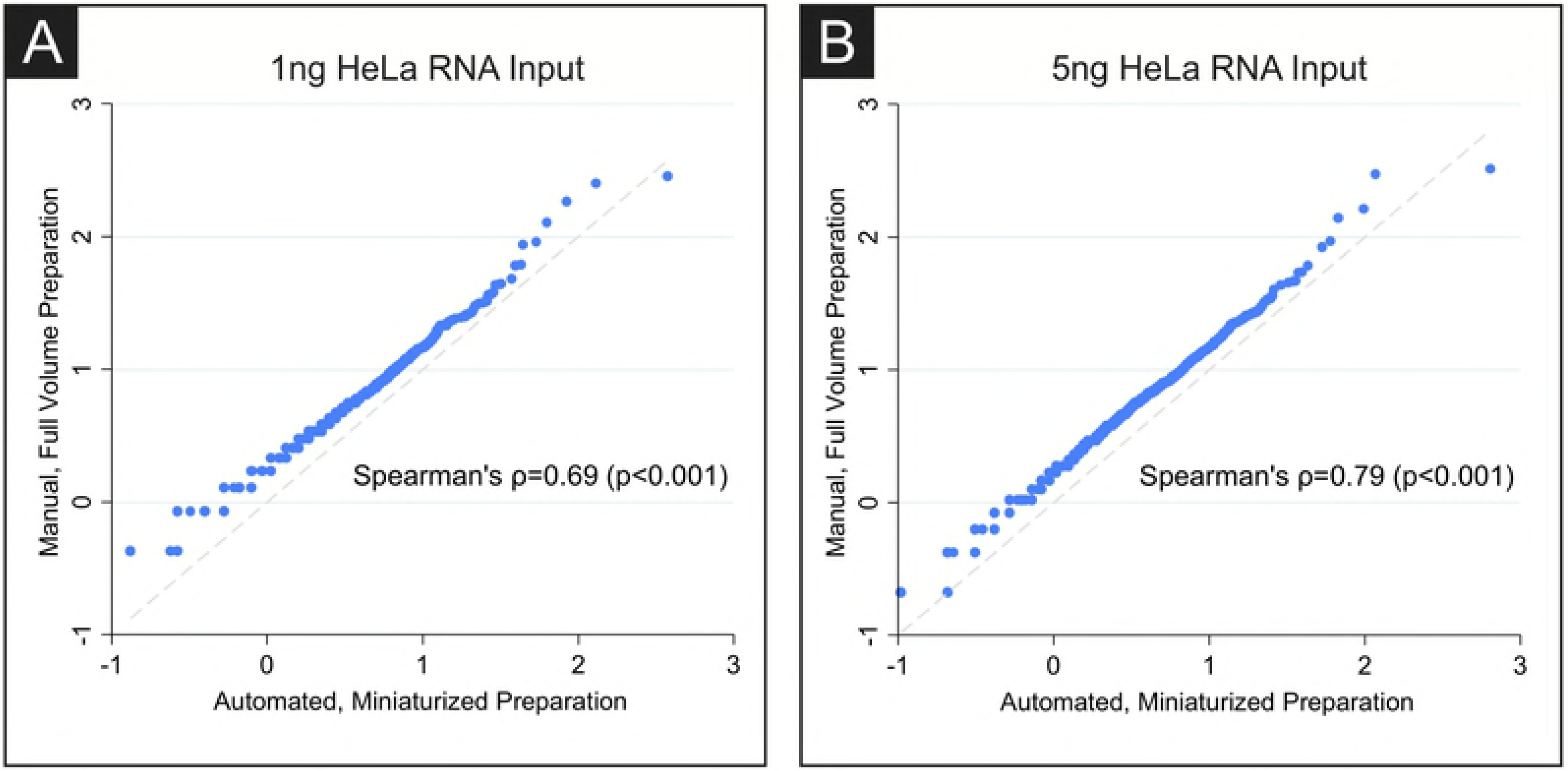
HeLa transcriptome coverage is comparable in full volume and miniaturized volume preparations. **Legend:** Rank-rank plots of the human transcriptome show strong correlation between the full-volume hand prepared protocol and the miniaturized, automated protocol for both (A) 1ng and (B) 5ng of HeLa RNA input (5ng RNA input: Spearman’s ρ = 0.79, p<0.001; 1ng RNA input: Spearman’s ρ = 0.69, p<0.001).

### Improving high-throughput library pooling

With advances in sequencing technology, the number of reads provided by a single run on high-throughput sequencers such as the HiSeq or NovaSeq is driving the creation of larger sample pools to take advantage of lower sequencing costs. Samples can be pooled to occupy equal portions of a flow cell lane by capillary electrophoresis, fluorimetry, or qPCR. These processes are costly, tedious, and error-prone due to imprecise estimations and inaccurate pipetting, especially when pooling large numbers of libraries together.

To overcome these difficulties, we employed a two-step process to optimize the pooling of hundreds of samples (**Fig 5**). First, we used the Echo to dispense *equal volumes* (500nL) from each sample of a set of 265 libraries. This pool was then sequenced on the Illumina iSeq to a total combined depth of 4.5 million reads. Use of the Echo to dispense nanoliter volumes allowed preservation of the bulk of each original sample library. The read distribution of the 265 libraries resulted in a normal distribution with each library occupying a mean of 0.377% ± 0.125% of the total reads per sample (**Fig 5**).

**Fig 5.**
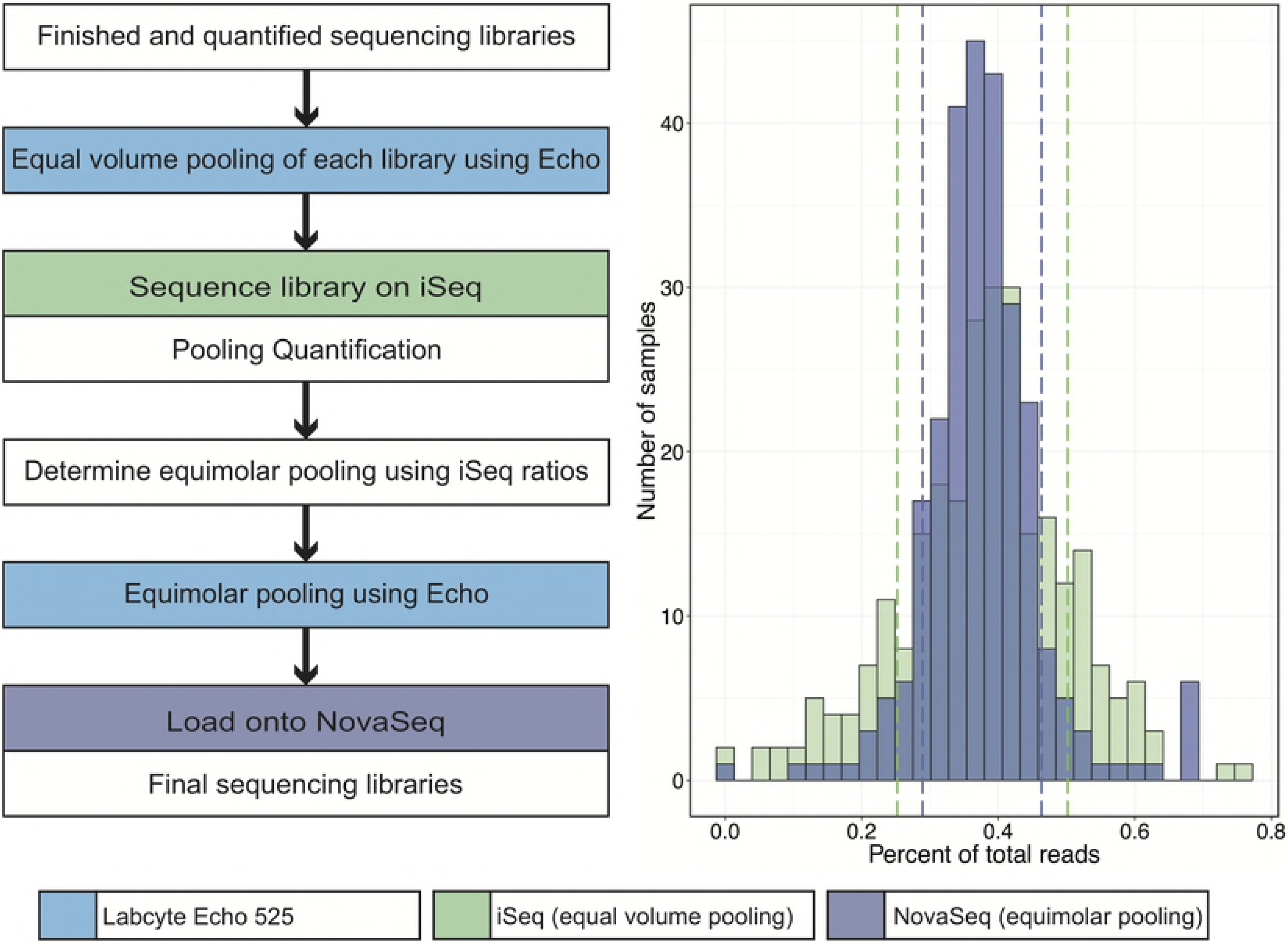
Pooling by iSeq correlates with pooling by NovaSeq. **Legend:** 265 libraries were pooled at equal volumes (0.5uL per library) using the Echo 525 and sequenced on the Illumina iSeq to a total combined depth of approximately 4.5 million reads (standard deviation indicated by green dashed lines). Resultant reads were assigned to each barcoded library to calculate the percent of total reads occupied by each sample. These ratios were used to re-pool each library in appropriate volumes (ranging from 160nL to 3800nL) using the Echo 525 to achieve equal read representation across all 265 samples. Libraries pooled at equal volumes and sequenced on the iSeq occupied a mean of 0.378% ± 0.125% of total sequencing reads (standard deviation indicated by purple dashed lines). Re-pooled based on iSeq ratios, libraries sequenced on the Illumina NovaSeq demonstrated a significantly tighter standard deviation, with each library occupying a mean of 0.376% ± 0.087% of total sequencing reads.

By virtue of the internal ERCC controls, the number of reads belonging to each library was proportional to the partial concentration occupied by each library. This enabled estimation of the partial concentrations for each of the original 265 libraries. Next, the Echo 525 was used to dispense calculated *equimolar* volumes of each library (ranging from 160nL to 3800nL) into a final pool. The pool was sequenced on the Illumina NovaSeq to a total combined depth of approximately 11 billion paired-end reads. Sequencing results from the NovaSeq yielded a mean library proportion of 0.376% ± 0.087% of the total reads per sample (in this case, 41.7 million ± 9.6 million paired-end reads per sample). The significantly tighter standard deviation produced by this step and shown in **Figure 5** demonstrate that hundreds of libraries can be pooled quickly and within close range of each other using this method. As expected, samples with low read counts on the original iSeq calibration run possessed the greatest variability when pooled for sequencing on the NovaSeq.

### Cost and time comparison

This miniaturized, high-throughput protocol significantly reduces the cost and time associated with library prep. The materials cost for each library preparation using the manual protocol, including consumables such as reagents and tips, was approximately $43 (~$16,648 for 384 samples). Using this miniaturized protocol, the cost per sample dropped to approximately $8 per sample (~$3,161 for 384 samples), resulting in cost savings of over 80% (**Table 3A)**. Similarly, the automation of this library prep resulted in significant time savings. To complete 384 samples by hand with the manual protocol would have consumed an estimated 166 hours (assuming 16 samples are prepared at a time); whereas the time to complete the same number using the automated miniaturized protocol was approximately 10 hours (**Table 3B**). This significant time savings further increases the cost savings of this protocol, as the cost estimates above do not take labor into account.

**Table 3A).** Cost comparison of full-volume and miniaturized protocols for 384 samples.

|  | Full volume protocol | Miniaturized protocol |
| --- | --- | --- |
| Reagents | \$13,427 | \$1824 |
| Ampure beads | \$2024 | \$282 |
| Tips | \$1085 | \$1020 |
| Plates/tubes | \$112 | \$14 |
| Source plate | N/A | \$21 |
| **TOTAL** | **\$16,648** | **\$3,161** |

**Legend:** Approximate cost comparison between the full volume protocol (done by hand) and the miniaturized, automated protocol (performed using the Gene-Vac EZ-2, Labcyte Echo 525, and the Beckman Coulter Biomek NX^P^). The miniaturized, automated protocol costs approximately 19% of the regular full volume hand prep. When accounting for employee salary for the time required to complete 384 libraries, the cost of the miniaturized protocol drops significantly.

**Table 3B).** Comparison of time at the bench for full-volume and miniaturized protocols for 384 samples.

|  | Full volume protocol | Miniaturized, automated protocol |
| --- | --- | --- |
| Calculations/planning | 120 minutes | 10 minutes |
| Aliquoting RNA | 120 minutes | 120 minutes |
| Drying RNA | N/A | 25 minutes |
| Master mix preparation | 120 minutes | 10 minutes |
| Dispensing master mixes | 215 minutes | 45 minutes |
| Incubations | 5808 minutes | 242 minutes |
| Bead cleans | 2880 minutes | 120 minutes |
| Quality check | Using the fragment analyzer and iSeq, QC time is equivalent for 384 samples |  |
| Pooling | 720 minutes | 20 minutes |
| **TOTAL** | **9,883 minutes (~166 hours)** | **592 minutes (~10 hours)** |

**Legend:** Approximate bench-time comparison between the full volume protocol (done 16 at a time by hand) and the miniaturized, automated protocol (performed using the Gene-Vac EZ-2, Labcyte Echo 525, and the Beckman Coulter Biomek NX^P^). The miniaturized, automated protocol can complete 384 libraries in approximately 6% of the time it would take to complete the same number by hand using the regular protocol.

### Use-case example: mNGS for pathogen detection

As a demonstration of the applicability of this protocol, we used mNGS for pathogen detection in clinical samples. Sequencing libraries were prepared, as described above, from RNA isolated from two clinical samples. **Patient #1:** Endotracheal tube aspirate from a patient with respiratory failure was collected with IRB approval and consent. An aliquot was sent to the UCSF hospital clinical lab and another aliquot was placed immediately on dry ice. Standard bacterial culture of this sample produced heavy *Enterobacter cloacae* growth. Sequencing libraries produced by hand with full reagent volume, and again as described in the miniaturized protocol above, both demonstrated robust detection of *E. cloacae* RNA (109 unique rpm vs 103 unique rpm, respectively). **Patient #2:** bronchoalveolar lavage sample was collected from another patient with respiratory failure, with IRB approval and consent. Standard bacterial culture of this sample produced heavy *Haemophilus influenzae* growth. Sequencing libraries produced concurrently by hand with full reagent volume, and as described in the miniaturized protocol above, demonstrated comparable detection of *H. influenzae* (160 unique rpm vs 126 unique rpm, respectively) in both preparations. Human-stripped FASTQ files are available at BioProject Accession #PRJNA493096.

## Discussion

In this study, we present a miniaturized, highly automated, high-throughput protocol for preparation of high-quality mNGS sequencing libraries. Compared to the standard full-volume protocol, this protocol is faster, less expensive, produces data of equivalent quality for pathogen detection and human transcriptome coverage in HeLa preparations, and, for use-case patient samples, demonstrates correlation with clinical microbiology test results. To our knowledge, this is the first application of the Echo for metagenomic and metatranscriptomic sequencing analysis and adds to existing Echo-based protocols for DNA synthesis and plasmid sequencing [20,21].

This protocol has several advantages over alternative library preparation approaches. First, since the Labcyte Echo does not use pipette tips, tip contamination is eliminated as is the pipetting error inherent to transferring small volumes by volumetric displacement [22]. Second, simultaneous processing of up to 384 samples significantly reduces variance between library preps and exposes all samples to the same environmental and reagent contaminants, minimizing batch effects. Third, use of ERCC spike-in controls allows the determination of the original RNA input quantity, which is highly variable in human patient samples, and is often difficult to measure by traditional spectroscopy. The ERRC controls also help establish the degree of linearity between read counts and input concentrations. The latter is particularly important to avoid calling false negatives in clinical metagenomics. Lastly, the robots used in this miniaturized and automated library prep protocol enable rapid and simultaneous processing of a up to 384 samples at a time. This results in significant cost and time savings and can allow large-scale projects to be completed at an expedited timeline.

There are several limitations to this protocol. First, although this protocol reduces the time needed to prepare libraries for sequencing, it does not take into account the necessary upstream procedures to isolate nucleic acid [23]. Second, Labcyte Echo source plates have a limited working volume which results in dead volume loss of reagents in each source well; however, because the overall volume of each master mix is greatly reduced by the miniaturization, this dead volume does not greatly affect the overall cost of the prep. Third, the bead-based clean-up steps described here require aspiration with traditional tip based liquid handlers. Finally, the capital expense cost of robots used in this protocol is significant. Access to a core lab facility with these machines will greatly reduce the initial start-up cost of this protocol.

In conclusion, we present an automated, miniaturized, high-throughput protocol to prepare RNA sequencing libraries using the NEB Ultra II RNA Library Prep Kit. With this workflow, it is possible to prepare 384 high-quality sequencing libraries with just 10% of the regular reagent volume, at less than 20% of the cost and in less than 10% of the time compared to the regular hand-prep. The workflow presented here may support the further advancement of clinical metagenomics as well as large scale sequencing projects.

## Acknowledgements

We thank Valentina Garcia, Katrina Kalantar, Matt Laurie, and Michael Wilson for lab assistance, Anna Sellas, Rene Sit, and Derek Bogdanoff for sequencing help, and Howard Lee from Labcyte for Echo advice. We also thank the patients and families who contributed to the work performed here.

